# Identification of key genes and pathways associated with Crohn’s disease by bioinformatics analysis

**DOI:** 10.1101/543835

**Authors:** Zheng Wang, Jie Zhu, Lixian Ma

## Abstract

Crohn’s disease is a type of inflammatory bowel disease posing a significant threat to human health all over the world. Genome-wide gene expression profiles of mucosal colonic biopsies have provided some insight into the molecular mechanisms of Crohn’s disease. However, the exact pathogenesis is unclear. This study aimed to identify key genes and significant signaling pathways associated with Crohn’s disease by bioinformatics analysis. To identify key genes, an integrated analysis of gene expression signature was conducted using a robust rank aggregation approach. A total of 179 Crohn’s disease patients and 94 healthy controls from nine public microarray datasets were included. MMP1 and CLDN8 were two key genes screened from the differentially expressed genes. Connectivity Map predicted several small molecules as possible adjuvant drugs to treat CD. Besides, we used weighted gene co-expression network analysis to explore the co-expression modules associated with Crohn’s disease pathogenesis. Seven main functional modules were identified, of which black module showed the highest correlation with Crohn’s disease. The genes in black module mainly enriched in Interferon Signaling and defense response to virus. Blue module was another important module and enriched in several signaling pathways, including extracellular matrix organization, inflammatory response and blood vessel development. There were also several other meaningful functional modules which enriched in many biological processes. The present study identified a number of key genes and pathways correlated with Crohn’s disease and potential drugs to combat it, which might offer insights into Crohn’s disease pathogenesis and provide a clue to potential treatments.

## Introduction

Crohn’s disease (CD) is a chronic nonspecific inflammatory bowel disease (IBD), which may affect any region of the gastrointestinal tract intermittently with the terminal ileum and colon being the most common [1]. The incidence and prevalence of CD were highest in developed countries, with the highest annual incidence in North America (20.2 per 100,000) and Europe (12.7 per 100,000). Similarly, the highest worldwide prevalence of CD was found in Europe (322 per 100,000) and North America (319 per 100,000) [2]. Furthermore, the incidence of CD in the developing countries continues to increase. Regarding sexual distribution, there was no sex preponderance in adult CD [1]. About 12% of CD patients have a family history of IBD, with an even higher proportion of family cases in younger individuals [3]. Although the exact etiology of CD remains uncertain to date, there is evidence that it involves a complex interaction between the individual’s genetic susceptibility, external environment, intestinal microbial flora and the immune responses [4, 5]. Previous genome-wide association studies have reported numerous loci associated with CD, which included NOD2, ATG16L1 and IRGM. NOD2 was identified as the first susceptibility gene for CD [6]. NOD2, a cytosolic pattern recognition receptor for the bacterial proteoglycan fragment muramyl dipeptide, plays a significant role in maintaining immunological homeostasis in the intestine. NOD2 mutations are genetically correlated with an increased risk for the development of CD [5, 7]. Moreover, genetic analyses have identified autophagy genes ATG16L1 and IRGM genes as susceptibility genes for CD [8]. Together, these common susceptibility genes account for 13.1% of variance in disease liability in CD [9].

A large amount of gene expression datasets are currently freely available from the Gene Expression Omnibus (GEO) database. Due to the small sample sizes in a single dataset and discrepancies of the characteristics within multiple heterogeneous datasets, it is necessary to integrate those massive datasets through systems biology tools and finally receive the stable and credible results [10]. So far, comprehensive integrated analysis of gene expression datasets in CD is still missing. Therefore, we executed integrated analysis method by using robust rank aggregation (RRA) method to detect the consistently significant differentially expressed genes (DEGs) from various expression profiling datasets [11]. In addition, we performed weighted gene co-expression network analysis (WGCNA) to categorise those DEGs into a number of functional modules, and followed by functional and pathway enrichment analysis [12]. Connectivity Map (CMap) represents a useful bioinformatic technique for establishing the connections among genes, drugs and diseases [13, 14]. CMap has been applied to discover drug action mechanisms and disease pathogenesis, as well as to assist in the identification of novel potential therapeutics [15, 16]. The DEGs of CD and control intestinal tissues were used to query the CMap database, which includes more than 7000 expression profiles corresponding to 1309 drugs. In the current study, we used CMap to explore therapeutic drugs that could potentially be effective against CD.

Here, we screened the differentially expressed genes from the intestinal tissue microarray data of CD patients and healthy people by means of bioinformatics technology, and obtained the signal pathways related to the disease through enrichment analysis. Moreover, the potential therapeutic drugs for CD were explored via the CMap database. The study will help to understand the pathogenesis of CD and lay a theoretical foundation for molecular diagnosis and targeted therapy.

## Methods

### Data Collection and eligibility criteria

The Probe signal data from the key word “Crohn’s disease biopsie” were downloaded from Gene Expression Omnibus (GEO) database(http://www.ncbi.nlm.nih.gov/geo/). The selection criteria for this study were displayed as follows: (a) gene expression datasets which included gene microarray chip technology; (b) studies comparing gene expressions between active Crohn’s disease and normal controls tissue in human samples; (c) the number of samples in each gene expression profiling dataset was no less than 6; (d) Processed data or raw data can be used for reanalysis should be provided in these databases. Studies not meeting these selection criteria were not remained in the analysis.

### RRA analysis

Data preprocessing steps and statistical analysis were performed using R software(https://www.r-project.org/) and Bioconductor packages [17]. Microarray datasets were normalized using RMA [18]and differential expression analysis was implemented using the limma R package [19]. The statistical operations were performed using the Robust Rank Aggreg package of R statistical software [11]. Genes were ranked by statistical significance which was evaluated by the expression fold change level and P values.

### Screening of small drug molecules

To explore potential therapeutic drugs against CD, the DEGs were compare to the expression signatures of 1309 drugs in the CMap database (https://portals.broadinstitute.org/cmap/) [13, 14]. The upregulated and downregulated DEGs generated through the RRA method that had a |logFC| >0.5 were imported into the CMap database for analysis. Compounds with negative connectivity scores, which indicated that the corresponding drug may reverse the expression of the query signature in CD, were recorded as potential therapeutic drugs. The small drug molecules with an average connectivity scores <-0.25 and P value <0.05 were recorded(Table 2).

### WGCNA

Gene co-expression network was implemented by the WGCNA package in R [12]. GSE16879 which including 37 CD patients and 12 controls was used to construct the WGCNA analysis. To include a sufficient amount of genes into WGCNA, genes with p<0.05 and |logarithmic fold changes| (|logFC|) >0.14 were chosen from the ranked gene list. When the soft threshold was defined as 11, the scale-free topology index was >0.9. In the present study, we chose a minimum module size of 150 for the genes dendrogram and the minimum height for merging modules at 0.15. The modules were randomly color-labeled.

### Functional enrichment analysis

Meaningful co-expression modules were screened out for functional enrichment analysis, performing by metascape (http://metascape.org) [20]. Top ten clusters with their representative enriched terms were selected.

### Statistical analysis

Statistical analyses were performed using R software version 3.5.0. P < 0.05 was considered to be statistically significant.

## Results

### CD microarray datasets

According to the screening criteria, 9 datasets in CD were finally included in this study: GSE10714 [21], GSE16879 [22], GSE36807 [23], GSE9686 [24], GSE10616 [25], GSE75214 [26], GSE6731 [27], GSE52746 [28], GSE68570 [29] (Table1). The description of the studies was provided in Table 1, such as GSE number, authors and number of samples. The number of CD patients in these studies ranged from 4 to 59, and the number of controls ranged from 3 to 22. A total of 179 CD patients and 94 healthy controls were finally included in this study.

**Table 1.**
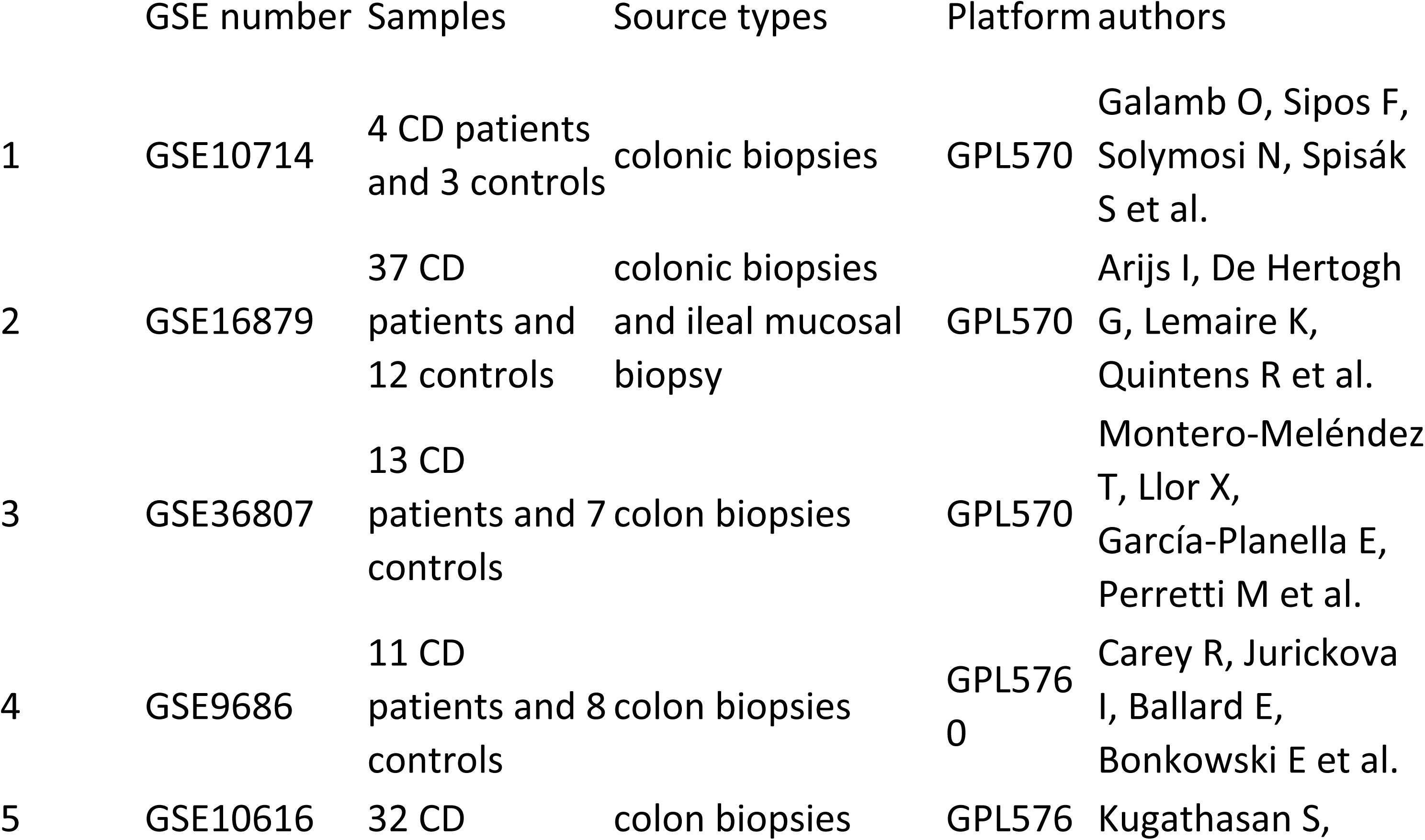
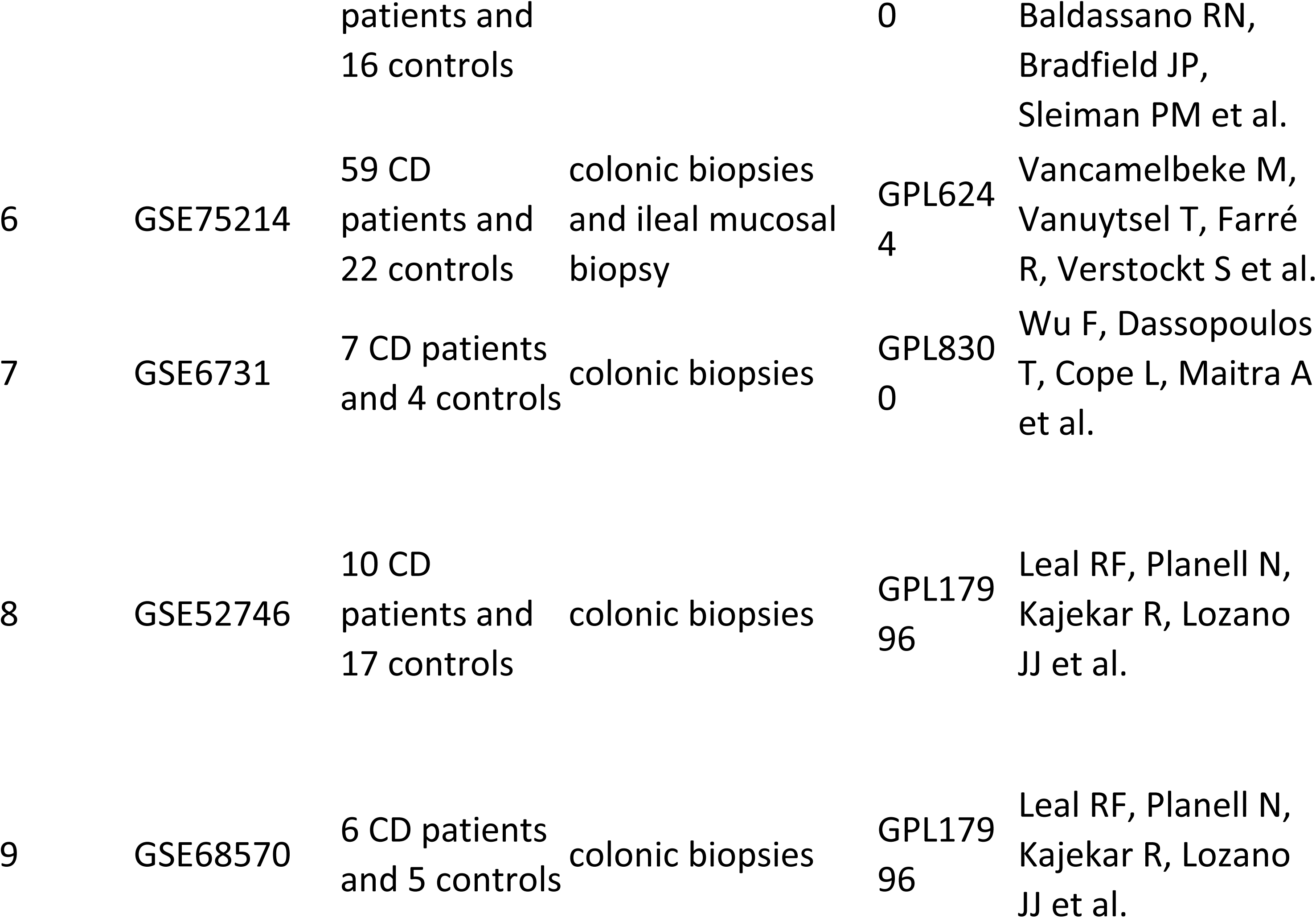
Summary of those 9 genome-wide gene expression datasets involving CD patients.

### RRA analysis

By using a RRA method to integrate 9 datasets, upregulated and downregulated ranked gene lists were obtained. S1 Table showed those top 150 upregulated and top 23 downregulated genes in CD patients. These top 150 upregulated genes had a p<7.4E −8 and a |logFC| >1. These top 23 downregulated genes had a p<2.1E −10 and a |logFC| >1. The top 20 upregulated and downregulated genes in the integrated microarray analysis were showed in Fig 1. Among these top genes, the abnormal expression of some genes had been well confirmed, such as MMP1, CLDN8 [30, 31]. However, the roles of other genes in CD have not been previously reported in the literature, such as PCK1, CHP2.

**Fig 1.**
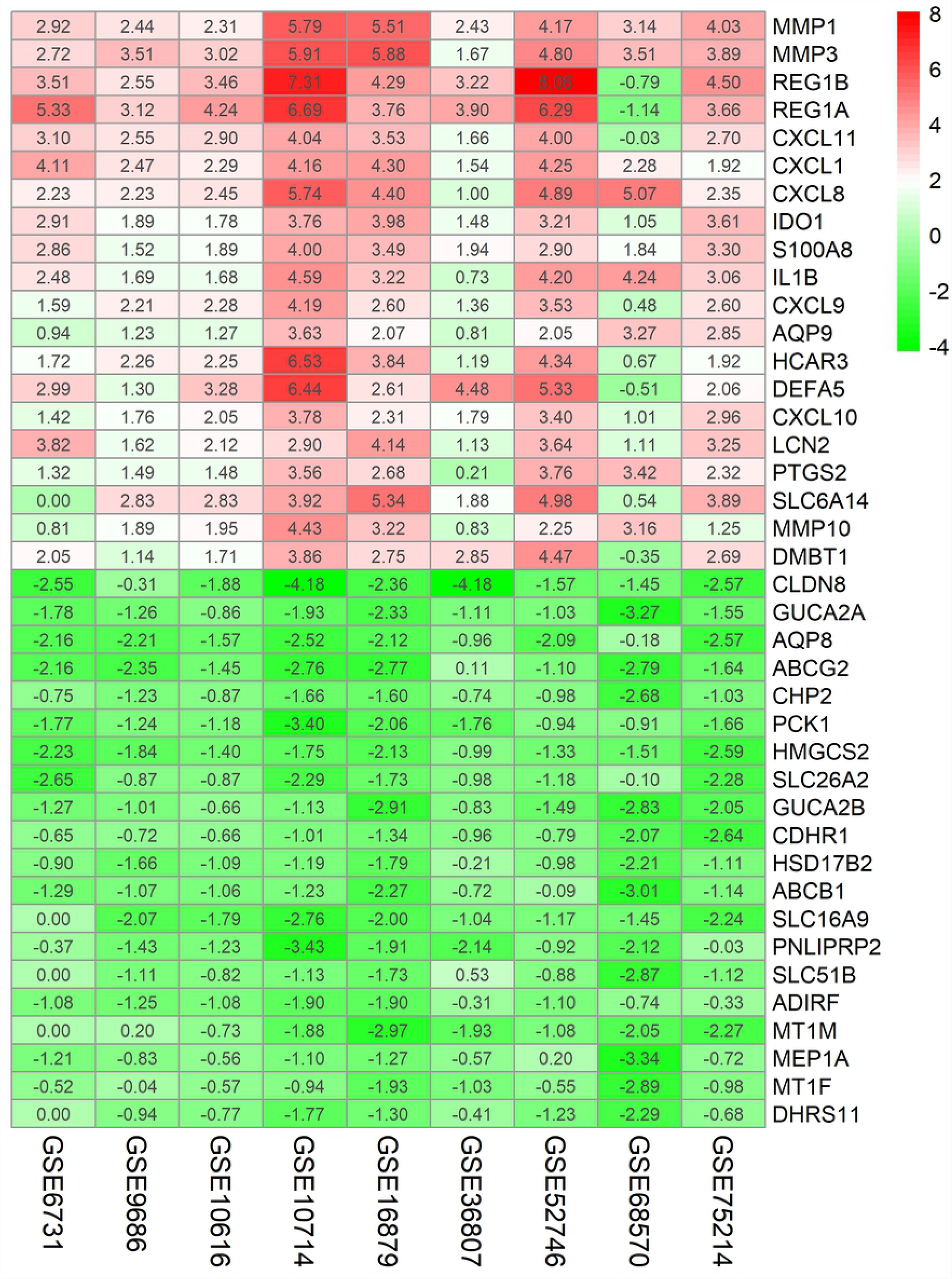
Top 20 upregulated and top 20 downregulated genes in CD. Each column represents one dataset and each row represents one gene. Numbers in each rectangle show the logarithmic fold change of genes in each dataset. Green indicates decreased gene expression and red indicates increased gene expression.

### Small drug molecule screening

The related small molecules with highly significant correlations were listed in Table 2. Among these molecules, tetracycline showed minimum p value and higher negative correlation, indicating the potential molecular signature that may counteract those of CD.

**Table 2.** Results of CMap analysis.

| cmap name | mean | enrichment<br>t | p |
| --- | --- | --- | --- |
| tetracycline | -0.41 | -0.9 | 0.00004 |
| methoxamine | -0.282 | -0.835 | 0.00135 |
| tubocurarine<br>chloride | -0.542 | -0.83 | 0.00151 |
| Prestwick-675 | -0.265 | -0.818 | 0.00203 |
| benzbromarone | -0.301 | -0.874 | 0.00403 |
| methocarbamol | -0.28 | -0.831 | 0.00959 |
| colistin | -0.266 | -0.724 | 0.0118 |
| disopyramide | -0.348 | -0.714 | 0.01361 |
| amantadine | -0.349 | -0.682 | 0.02222 |
| pepstatin | -0.274 | -0.638 | 0.04144 |

### WGCNA

The expressions of 5666 DEGs generated from the RRA analysis were used for WGCNA, all of which had a p<0.05 and a |logFC| of >0.14. As shown in Fig 2A, a power value of 11 was selected as the soft-thresholding to ensure a scale-free network. We set 150 as the least gene number of each gene network and 0.15 as the threshold for the merge of similar module. The WGCNA elucidated 7 coexpressed modules ranging in size from 269 to 1730 and the grey module represented a gene set that was not assigned to any of the modules (Fig 2B). Interactions of the 7 co-expression modules were analyzed (Fig 2C). The genes in each module were listed in S2 Table.

**Fig 2.**
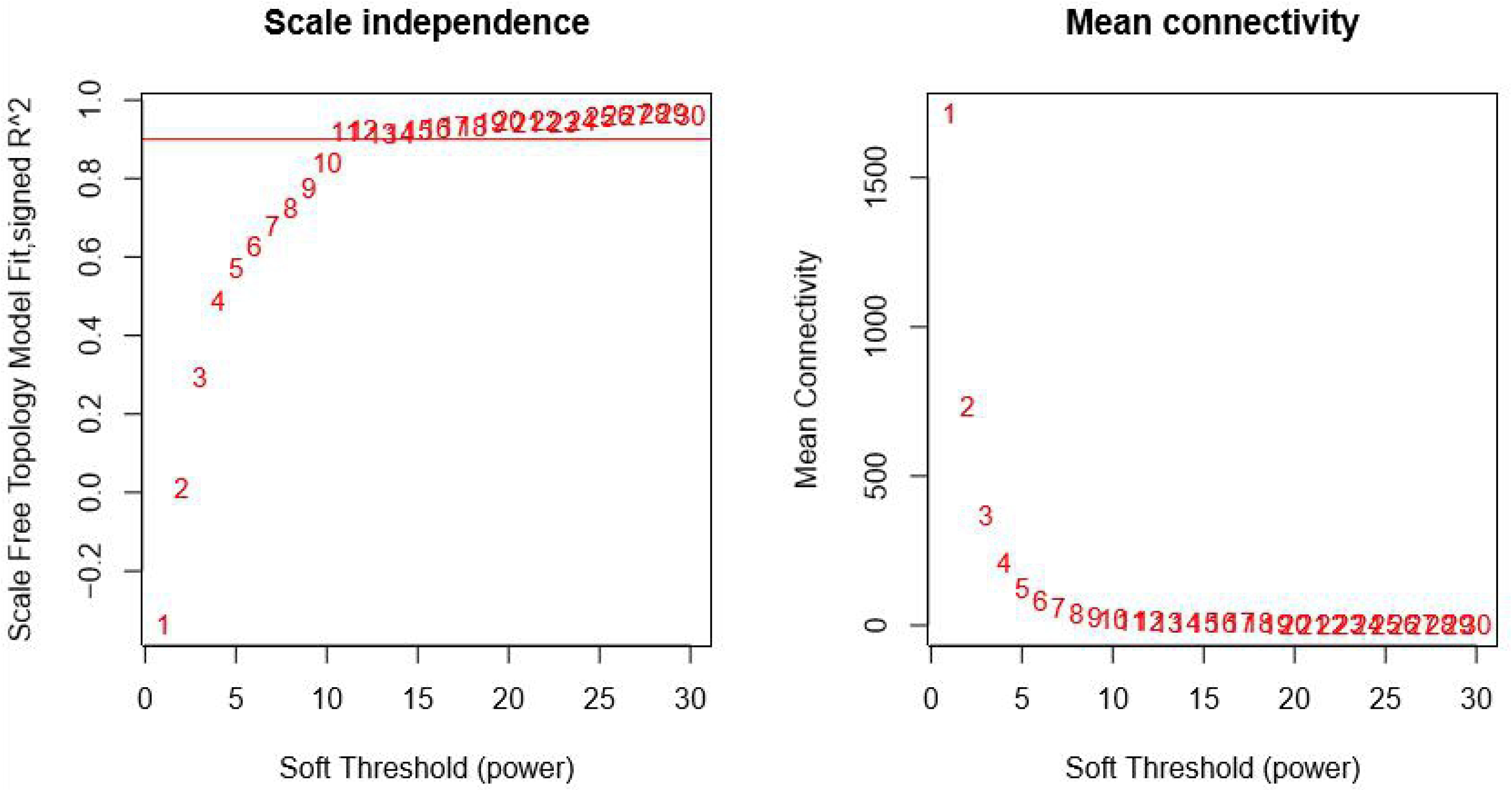
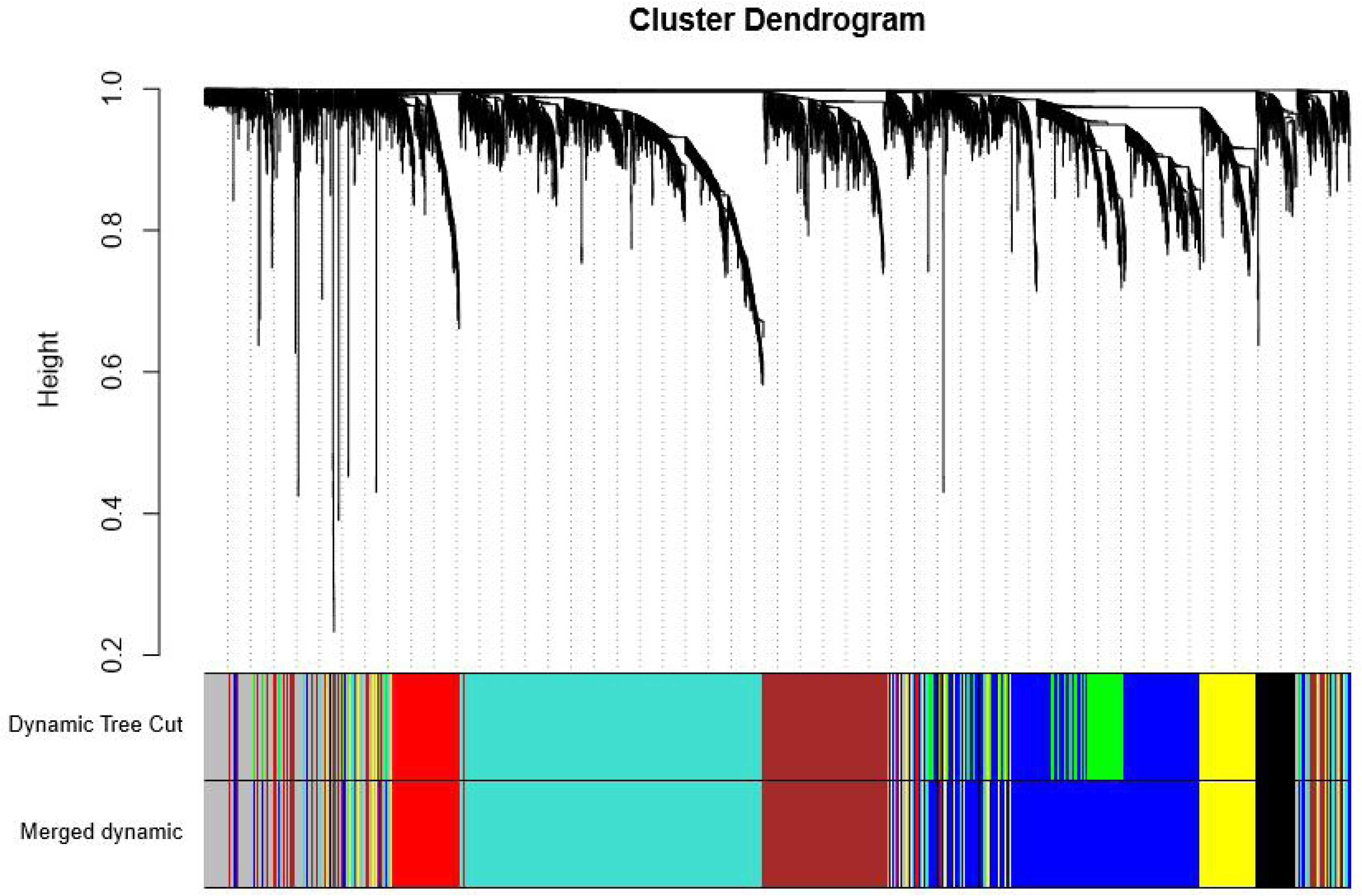
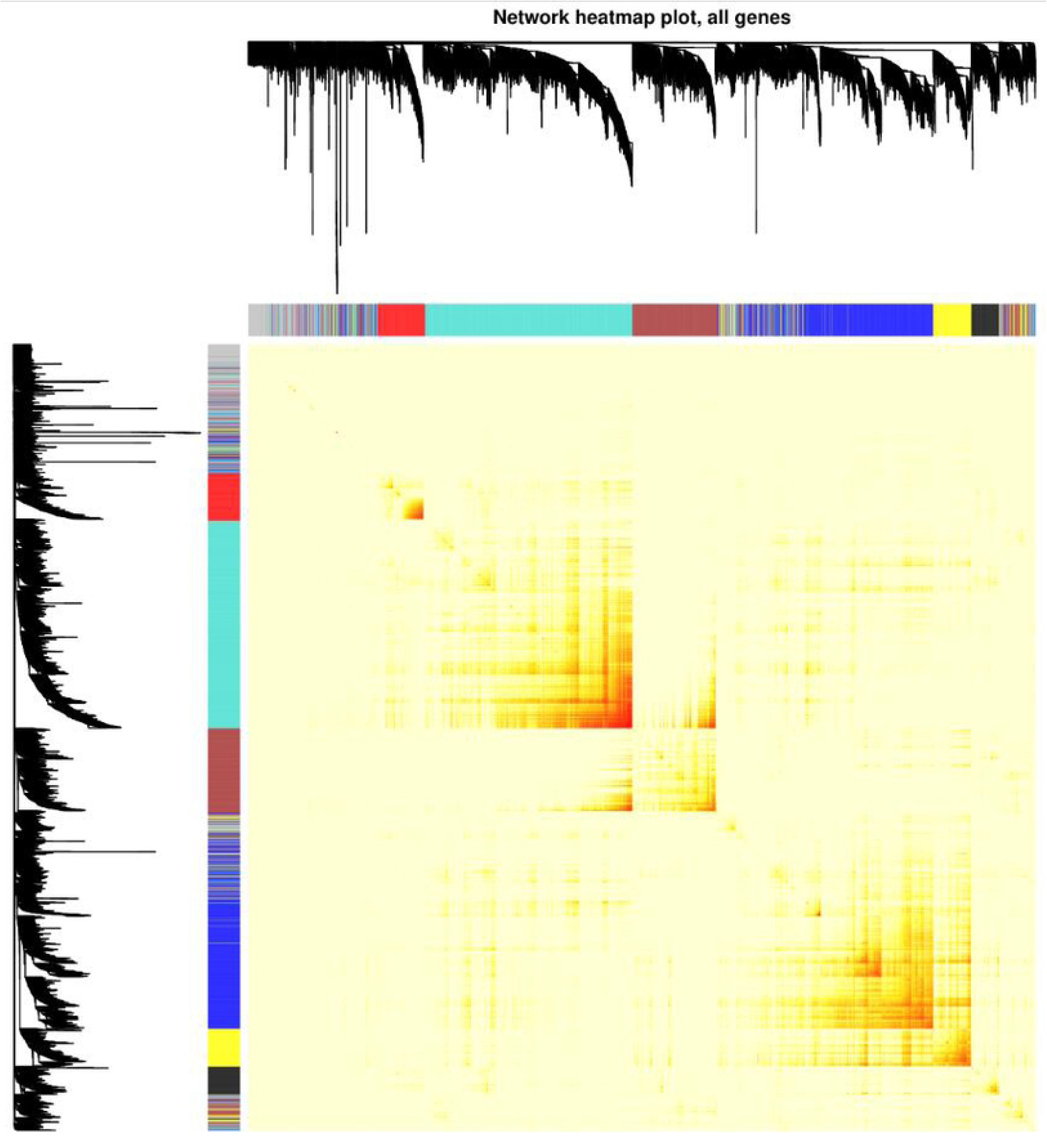
Plots in the WGCNA analysis using gene expressions in 37 CD patients and 12 controls from GSE16879 datasets. (A) Network topology of different soft-thresholding powers. The left panel displayed the influence of soft-threshold power on scale-free topology fit index; the right panel showed the influence of soft-threshold power on the mean connectivity. (B) Cluster dendrogram of coexpression genes and functional modules in CD. 7 co-expression modules were constructed and was shown in different color. (C) Visualizing the gene network using a heatmap plot. Different colors of horizontal axis and vertical axis represented different modules. Light color represents low overlap and gradually dark red represents higher overlap.

Identifying the modules most correlated with clinical features has a great biological significance. As shown in Fig 3, black, blue, red and yellow modules positively correlated to CD (p < 0.05). Turquoise and brown modules negatively correlated to CD(p < 0.05). Brown and grey modules were positively correlated with a better clinical response to infliximab (p<0.05) while blue and yellow modules was correlated negatively to treatment. Combined with Fig 4,we found that these six modules yielded two main clusters; one contained four modules (black, blue, red and yellow module) while the other contained two modules (brown and turquoise module). Moreover, two pairs of module combination had much higher adjacency degree and they are blue module and yellow module, black module and red module respectively. The black module showed the highest correlation with CD (r = 0.71, p = 1e-8, Fig 3). The genes in black module mainly enriched in Interferon Signaling and defense response to virus. Genes in red module were enriched in Cell Cycle, cell division and cell cycle phase transition. Genes in blue module were enriched in several signaling pathways, including extracellular matrix organization, inflammatory response, blood vessel development and leukocyte migration. The yellow module was similar to the blue module, mainly enriched lymphocyte activation, adaptive immune response, leukocyte activation involved in immune response. Some other signaling pathways were observed to enrich in other modules, such as monocarboxylic acid metabolic process, cofactor metabolic process and Metabolism of lipids in turquoise, Transport of small molecules in yellow. The main findings for the functional enrichment analysis were shown in Table 3.

**Table 3.**
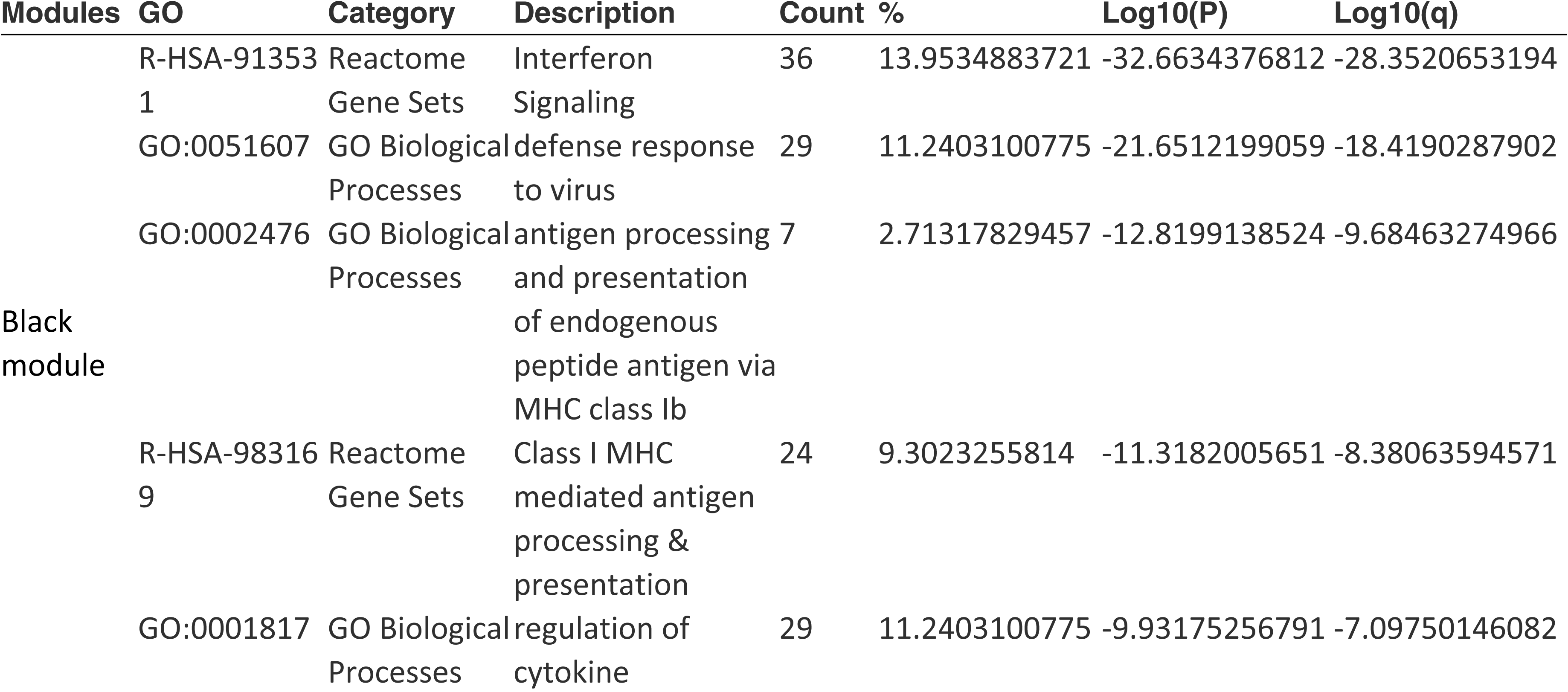
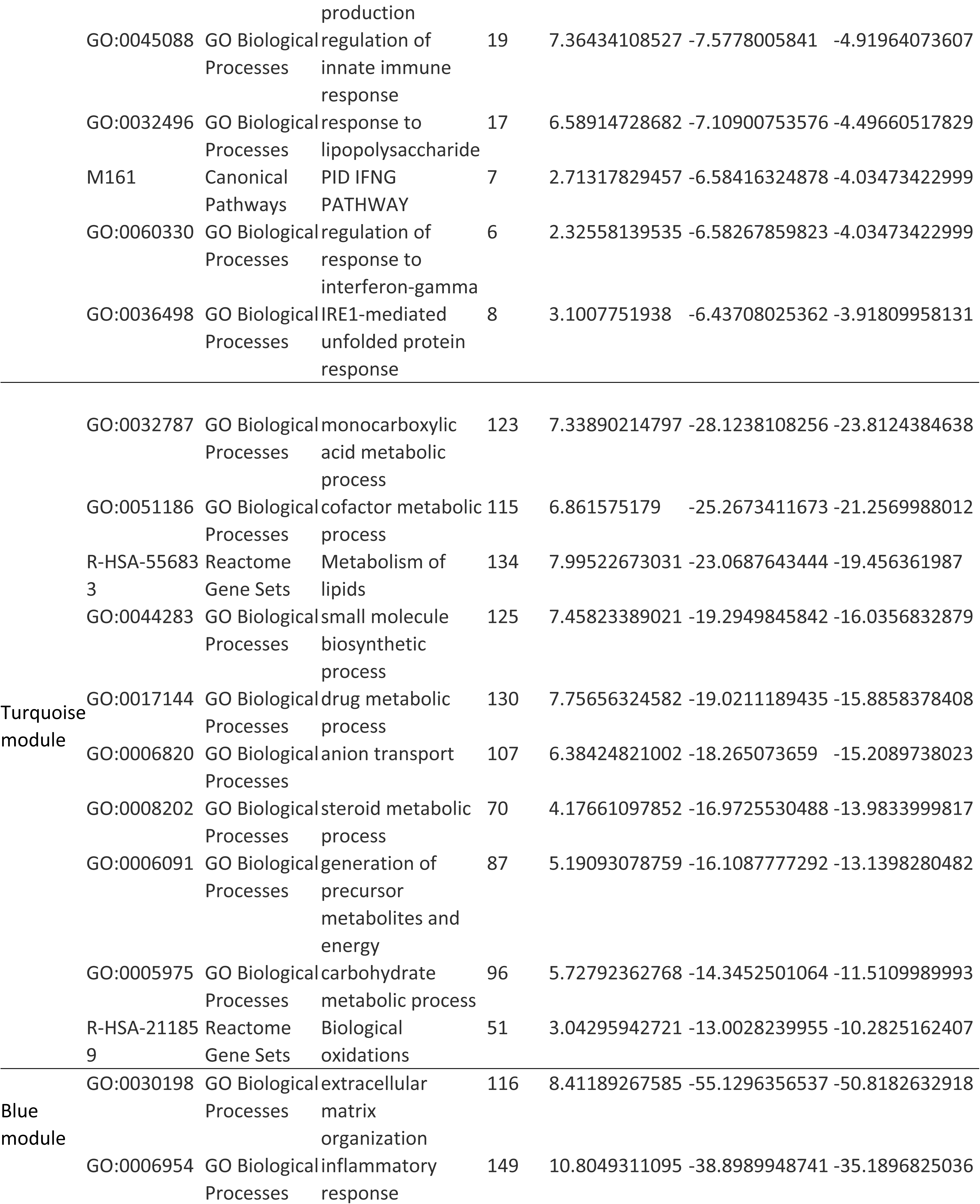
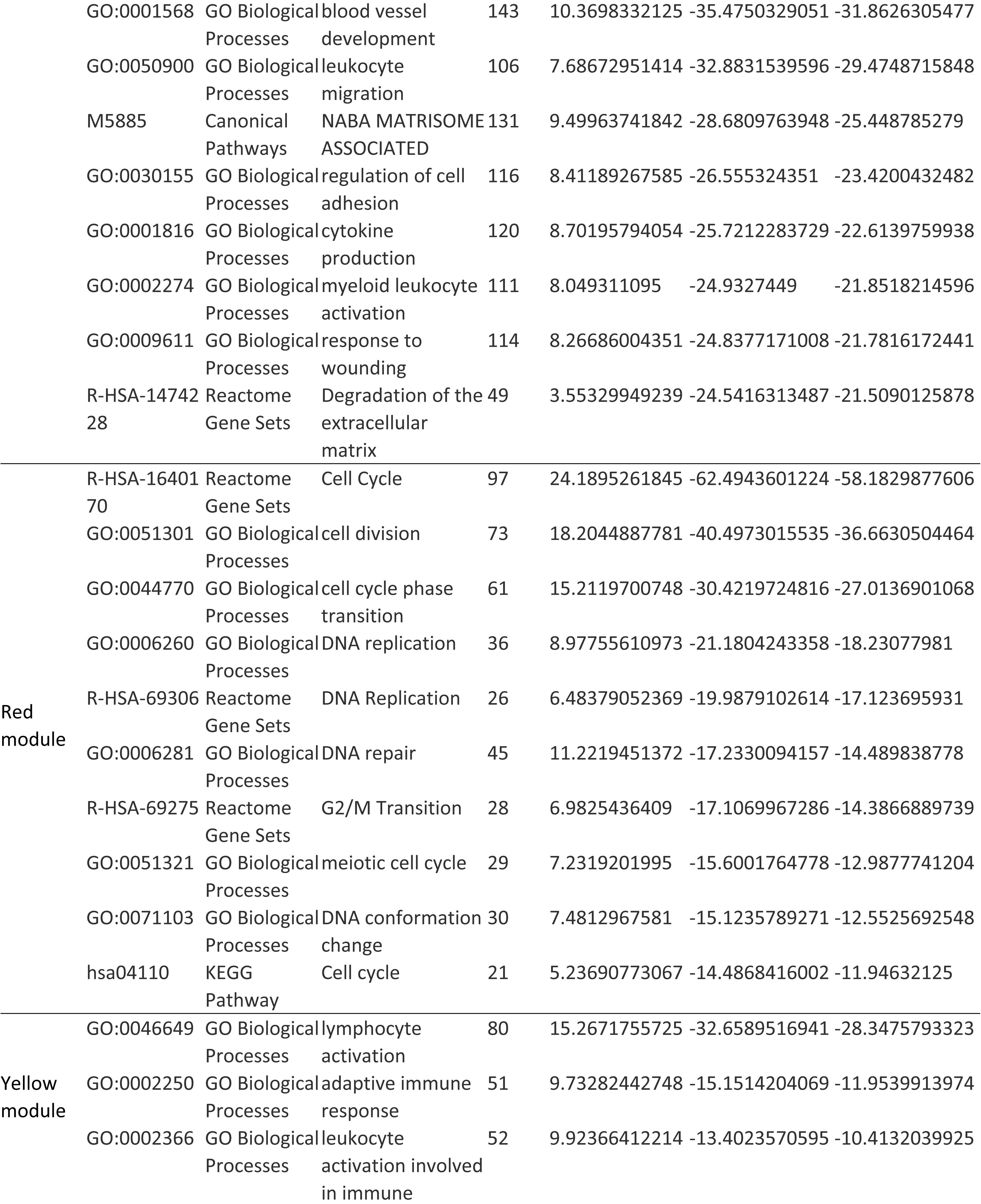
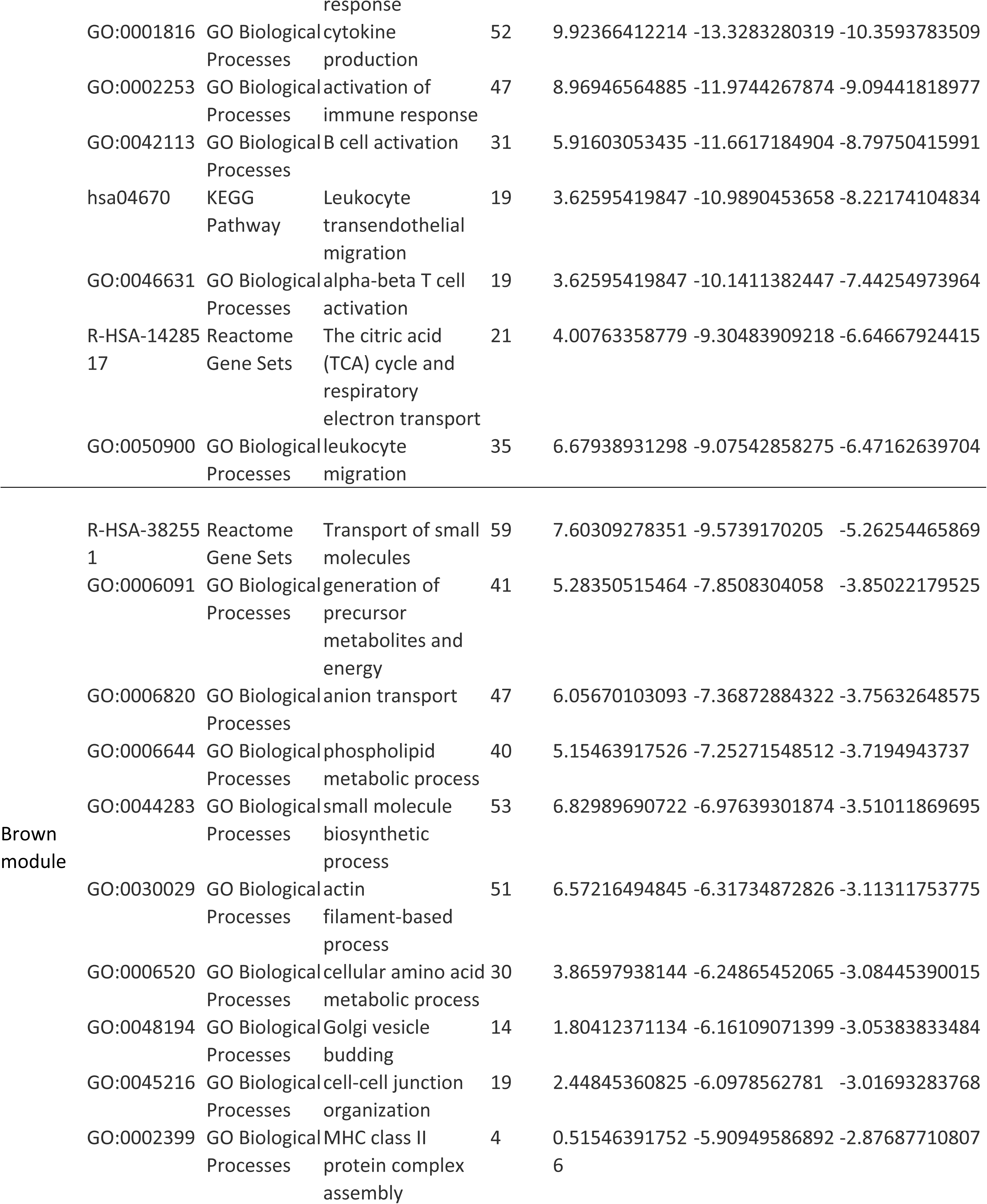
Pathway and process enrichment analysis of those functional coexpression modules in CD.

**Fig 3.**
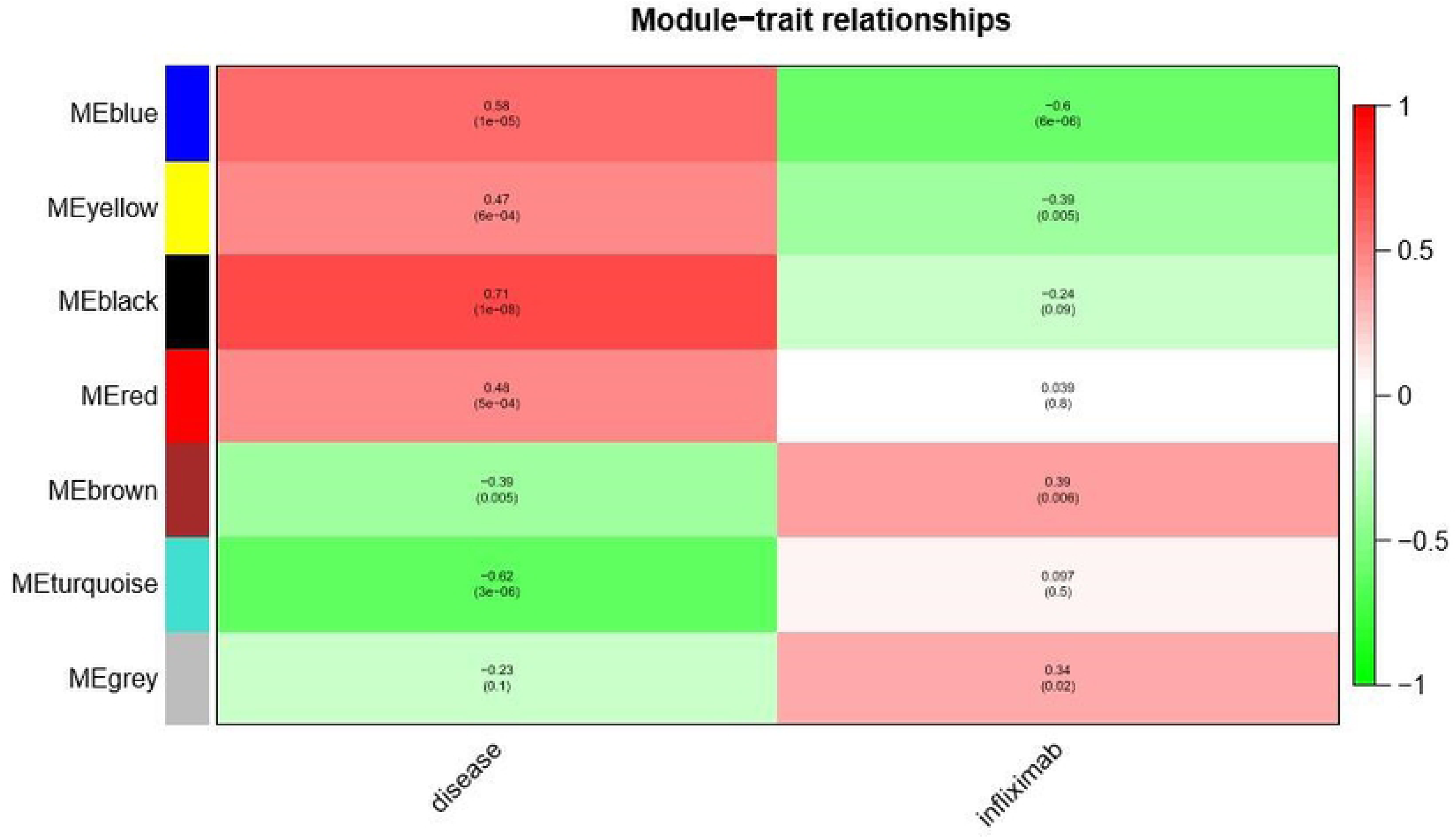
Heatmap of the correlation between module eigengenes and clinical traits of CD. The table is color coded by correlation according to the The table was color coded by correlation according to the color legend Each cell contains the corresponding correlation and p-value

**Fig 4.**
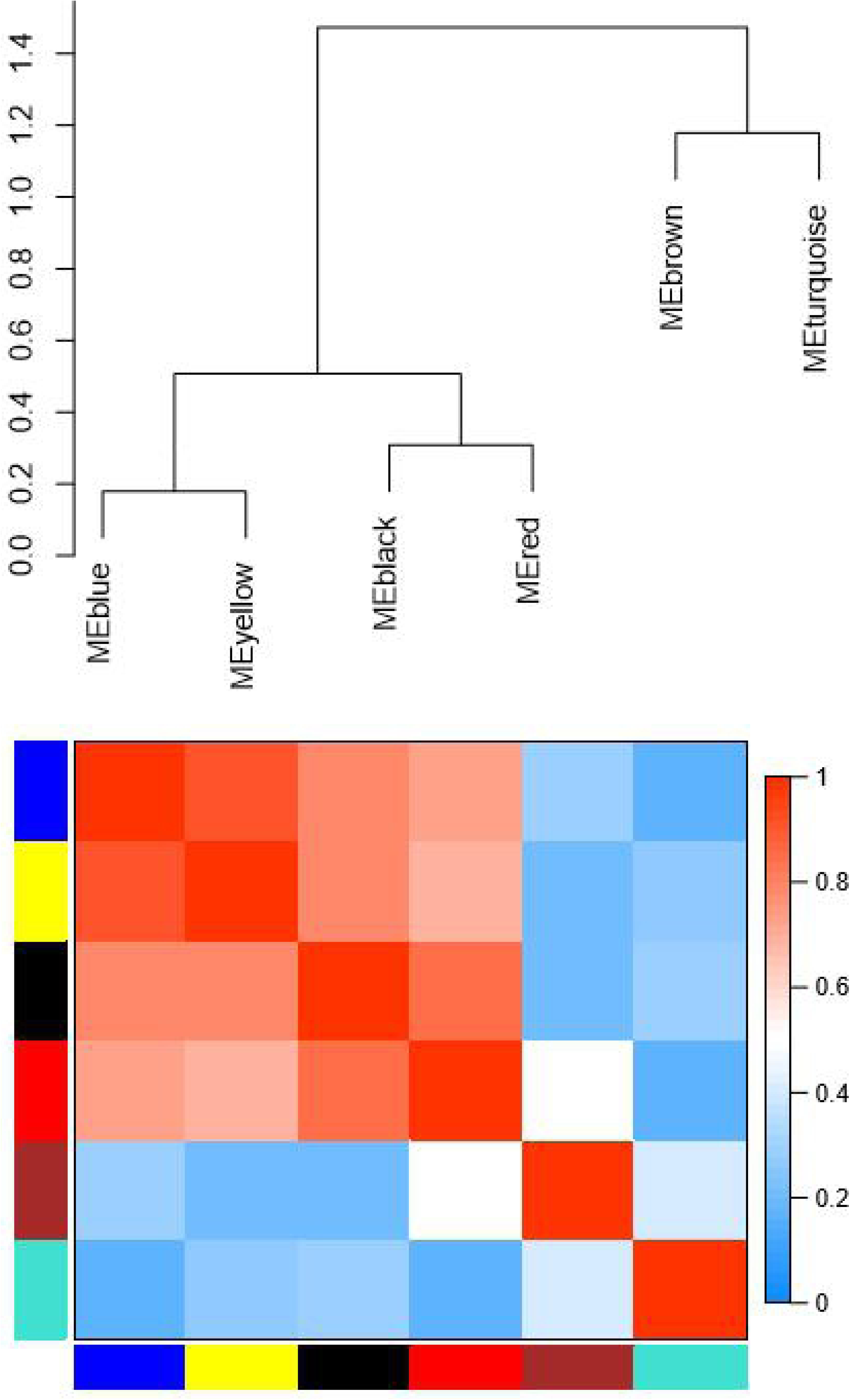
Heatmap plot of the adjacencies of modules. The change of color from blue (0) to red (1) in the heatmap represented the connectivity degree of different modules from weak to strong.

## Discussion

Crohn’s disease is a chronic inflammatory disorder which usually Lead to pathological changes at different locations along the length of gastrointestinal tract. The precise cause of CD has not been determined. In this study, we integrated the gene expression data to shed light on the pathological mechanism of CD.

A large number of remarkably upregulated and downregulated genes were identified through RRA analysis, some of which have been reported in previous literature in CD, such as MMP1, MMP3, REG1B, REG1A, CLDN8, GUCA2A. Previous studies have reported that the expression of MMP1 and MMP3 were upregulated in the colonic mucosa with CD compared with control [32]. MMP1 and MMP3 belong to the family of metal dependent enzymes, which are capable of degrading a wide range of extracellular matrix components [33]. It is of note that MMP1 was involved in inflammation and have been implicated in tissue injury and repairing processes in CD [34]. In accordance with previous reports, the present study found that MMP-1 and MMP-3 were located in the blue module, which was mainly enriched in extracellular matrix organization and inflammatory response. Because MMP-1 and MMP-3 are the top two of upregulated differentially expressed genes ranking, over expression of MMP-1 and MMP-3 are probably key factors in the pathogenesis of CD. It was reported previously that REG1B and REG1A were highly expressed in CD colonic mucosa [35]. However, the molecular mechanism underlying how REG1B and REG1A exert their role in CD has not been determined yet. CLDN8 is a member of the family of claudins, which mainly located in the cell membrane and associated with tight junctions of cell adhesion [36]. It has been reported that CLDN8 might play a significant role in the injury of intestinal epithelial barrier of CD [37]. The expression level of CLDN8 was remarkably downregulated in CD colonic mucosa compared to normal controls, but the molecular mechanism underlying its role remains unclear. As previously demonstrated, GUCA2A expression was markedly decreased in CD, but its molecular mechanism is poorly understood [38]. The other genes (such as PCK1,CHP2) are novel gene signatures of CD, and there is still lack of researches on its role in CD pathogenesis. Therefore, their aberrantly expressions need to be validated in future researches.

Several small molecules with potential therapeutic efficacy against CD were identified according to the CMap database. As a result of our screening effort, tetracycline has been identified as a potential therapeutic target for CD. Tetracycline were discovered in the 1940s and showed broad spectrum inhibitory activity against a wide range of microorganisms. Previous findings indicated that tetracycline exert pleiotropic functions independent of their antimicrobial activity, which include inhibition of MMPs, angiogenesis and inflammation [39]. Owing to the ability to inhibit MMPs expression and treat overgrowth of bacteria in the small intestine, tetracycline may be prescribed to treat CD. We might suppose that tetracycline could play certain roles to combat CD; however, further investigation on a large population is required to firmly establish the therapeutic effect in CD.

The WGCNA which is a kind of bioinformatics analysis method was conducted to explore the molecular mechanisms of various disease [40, 41]. Several reports involved IBD had been studied by this method [42, 43]. In this study, 5666 CD associated genes generated from RRA analysis were used in WGCNA and most of them were classified into 7 co-expression modules. Six of the modules have significant associations with disease (Fig 3), which can be enriched more refined in three cluster (Fig 4), highlighting some new insights into the pathogenesis of CD. To further understand the importance of those modules in the pathogenesis of CD, enrichment analysis was performed using metascape. The two most critical modules, black module and blue module, which have the most significant relationship with disease need to be addressed. Black module was mainly enriched in some biological processes, the most important of which were Interferon Signaling, defense response to virus (Table 3). It is generally known that IFN-gamma plays a key role in the early steps of installation of inflammation, promoting monocyte recruitment and activation, and inducing the expression of other inflammatory cytokines. Interferon Signaling has been identified as a central aspect of innate immune response which induces a wide variety of antiviral proteins against pathogens infection [44, 45]. Furthermore, previous studies have shown that Epstein-Barr virus and cytomegalovirus were the two main viruses that are positively correlated with the onset of CD [46]. In addition, bacteriophage was other viral agent that has been suspected to play a role in CD pathogenesis [47]. Although there was not enough evidence that the virus is directly involved in CD, the study by Cadwell et al. [48]suggested that the combination of host genetic susceptibility and the presence of viral factors could lead to CD occurrence. What’s more, several studies have found that for some CD patients with steroid resistance, antiviral therapy may be effective for relieving symptoms and improving the sensitivity of anti-inflammatory drugs [49]. From what have been discussed above, we may get the hypothesis that for some CD patients, especially those with steroid-refractory CD, virus infection and the sequential Interferon Signaling pathway may be the key factor in the initial stage of disease onset. Similarly, the enrichment analysis of genes in the salmon module mainly involved in extracellular matrix organization, inflammatory response and blood vessel development. Meanwhile, yellow module clustered with blue module mainly enriched in the pathway of lymphocyte activation. The pathways enriched from the two important modules were all belong to the intermediate link of the inflammatory response process, which explains the general pathogenesis of CD, an autoimmune disease. Moreover, extracellular matrix organization and blood vessel development also play a key role in the process of intestinal fibrosis at the late stage of CD [50]. Intestinal fibrosis continues to develop after intestinal primary injury and inflammatory response are subsided [51]. As we know, severe intestinal fibrosis is the main cause of intestinal resection, which seriously affects the prognosis of patients. In other words, extracellular matrix organization, inflammatory response and blood vessel development are key factors in the development of CD.

## Conclusion

In conclusion, this study identified a number of key genes and co-expression biologically functional modules which may be involved in the progress of CD, based on differentially expressed genes between CD samples and normal mucosas. Additionally, the present study screened a small drug molecule named tetracycline, which might be exploited as adjuvant drug for improved therapeutics for CD. Our findings suggest that virus infectious may be the initial link in CD. Moreover, extracellular matrix organization, inflammatory response and blood vessel development are the key pathways for the development of CD, which not only participate in the stage of inflammatory response process, but also play a key role in the late phase of intestinal fibrosis. This point may provide a new idea for improving the prognosis of CD.

## Acknowledgments

This work did not receive any specific funding from any sources.

## Author Contributions

Conceptualization: Zheng Wang, Jie Zhu, Lixian Ma

Data curation: Zheng Wang, Jie Zhu

Formal analysis: Zheng Wang, Jie Zhu, Lixian Ma

Investigation: Zheng Wang, Jie Zhu, Lixian Ma

Methodology: Zheng Wang

Software: Zheng Wang

Supervision: Lixian Ma

Visualization: Zheng Wang

Writing-original draft: Zheng Wang, Jie Zhu

Writing-review & editing: Zheng Wang, Jie Zhu, Lixian Ma

## Supporting information

**S1 Table.** Top 150 up-regulated and top 23 down-regulated genes in CD patients.

**S2 Table.** Six main functional co-expression modules and related genes involved in CD.

